# Characterization of Reconstructed Human Epidermis in a Chemically-defined, Animal Origin-free Cell Culture System

**DOI:** 10.1101/2024.02.13.580080

**Authors:** Julia Bajsert, Valerie De Glas, Emilie Faway, Catherine Lambert de Rouvroit, Miguel Perez-Aso, Paul W. Cook, Yves Poumay

## Abstract

The Reconstructed Human Epidermis (RHE) model derived from epidermal keratinocytes offers an ethical and scientifically valid alternative to animal experimentation, particularly in cutaneous toxicology and dermatological research, where elimination of animal cruelty is of paramount importance. Moreover, safer cell- and tissue-based cutaneous therapies are also possible, where the removal of animal-originated products from the RHE culture media lessens the risk of adverse clinical events. Thus, we compared chemically-define animal origin-free (cdAOF) supplements and the historically utilized supplement (HKGS), which contains growth factors and bovine pituitary extract. Herein we present the first extended characterization of RHE using cdAOF culture systems with newborn, adult, and immortalized N/TERT keratinocytes. Culture of RHE in the cdAOF media produced histological features that were nearly identical to that produced using HKGS, with the exception that the basal keratinocytes were less cylindrical, and with some immunolocalization of involucrin in the basal layer. Additionally, increased mRNA expression of several inflammatory-proliferative markers was observed in the cdAOF RHEs as well. Importantly, in RHEs cultured in cdAOF media, expression and immunolocalization of other expected markers of keratinization, as well as monitoring of barrier function revealed results equal or close to those observed in RHE cultured in HKGS.

## INTRODUCTION

The disadvantages of using human or non-human animal-originated products (e.g. serum, plasma, platelet lysates, serum-derived albumin, serum-derived transferrin, pituitary extract) or feeder layer cells (e.g. immortalized murine 3T3 fibroblasts, human dermal fibroblasts) in cell culture media have been emphasized since the inception of mammalian cell culture. For example, using animal-originated products can increase the expense of culturing mammalian cells, increase media inter-lot variability, and increase the risk of infection with known human pathogens that include Hepatitis, HIV, or mycoplasma, and transmissible spongiform encephalopathies (e.g. bovine spongiform encephalopathy/mad cow disease (BSE), chronic wasting disease (CWD), Creutzfeldt-Jakob (CJD)). Other known, or even yet to be characterized human and zoonotic infective agents might also pose risks to future cell- and tissue-based therapy. Because of these risks, increased regulatory pressure exists to develop chemically-define animal origin-free (cdAOF) cell culture systems for clinical applications. Collectively, the development and use of cdAOF cell culture supplements and media are thought to promote safer, pathogen-free cell cultures for cells, engineered human tissue, and engineered organs for human therapy.

Over the last several decades, research and development efforts have been made to reduce or eliminate animal-originated products from cell culture media used for normal non-transformed, non-immortalized cells. As an example, for human neonatal (HEKn) and adult (HEKa) epidermal keratinocyte cell culture, DMEM/F12, serum, cholera toxin and murine 3T3 feeder layer cells were replaced with improved modified MCDB 153 basal medium, or EpiLife medium, purified growth factors and animal originated products that included bovine pituitary extract (BPE), bovine serum-derived albumin (BSA) or bovine serum-derived transferrin (Boyce & Ham, 1983; Peehl & Ham, 1980; Rheinwald & Green, 1977; Rheinwald & Green, 1975; Shipley & Pittelkow, 1987; Tsao et al., 1982). Some of these epidermal keratinocyte cell culture systems were characterized as defined, serum-free or feeder layer-free, but not cdAOF. Eventually, a cdAOF human keratinocyte cell culture system, Supplement S7 (a.k.a. S-70203, S70203) containing growth factors, hormones and other proteins was reported in 2004 for use with EpiLife basal medium (Cook PW et al., 2004a). Using the cdAOF Supplement S7 - EpiLife system, it was shown that HEKa could be efficiently isolated in primary culture and serially propagated, with the requirement for pre-coating the cell culture substrate with recombinant human collagen-1. Importantly, HEKa reared in the Supplement S7 - EpiLife system were demonstrated to form a cdAOF reconstructed human epidermis (RHE) that was histologically similar to human epidermis, but with a less columnar basal epidermis, a compacted stratum corneum and basal cell layer expression of involucrin. Importantly, this work demonstrated the first example of an engineered mammalian tissue (RHE) generated from primary cell culture under complete cdAOF cell/tissue culture conditions.

More recently, an improved cdAOF cell culture supplement system (HKSdaFREE + HKGE in EpiLife basal medium) was developed that supports both the cdAOF primary isolation and serial propagation of HEKn and HEKa, as well as human corneal epithelial cells (Cook P et al., 2012; Cook PW et al., 2012). Compared to Supplement S7, post-primary propagation was not dependent on precoating the cell culture surface with recombinant human collagen-1. This new cdAOF HEKn culture system also supported the generation of RHE that was histologically similar to other RHE models as well as to normal human skin (Cook PW et al., 2012). Furthermore, to demonstrate the therapeutic potential under cdAOF cell culture conditions, HEKn isolated and propagated in EpiLife basal medium plus the HKSdaFREE + HKGE supplements were utilized as a novel cdAOF epidermal cell bioink to generate xenofree, fully vascularized full-thickness 3D-bioprinted skin equivalents that were successfully engrafted onto mice (Baltazar et al., 2023). Finally, using a related variant cdAOF supplement system (HFSdaFREE + HFGE + HFGE2), efficient primary and post-primary propagation of human dermal fibroblasts as well as corneal fibroblasts were demonstrated as well (Cook P et al., 2012; Cook PW et al., 2012). Collectively, these new cdAOF cell culture system supplements represented a breakthrough for the improved cdAOF culture of human skin- and cornea-derived cells, for potential use in cell- or engineered tissue-based therapies.

Since a ban on animal testing of cosmetic products was announced and then rendered efficient in 2013 by the European Union, it prompted the development of alternatives to previous tests that use live animals for skin studies. Additionally, reduction of animal cruelty by the elimination of animal-originated cell culture media components, such as fetal bovine serum (FBS), BPE, transferrin and BSA from keratinocyte and RHE culture media, has also been of interest. Within this context, models reconstructing human skin by cell culture were developed and characterized, before evaluation and acceptance for cosmetic testing were recognized under the current OECD guideline 431 (OECD, 2015). Protocols were therefore published to widely spread know-how of epidermal reconstruction and the Open-Source principle has been proposed and applied for collective contribution to potential improvements. Such models are of course now shown to be scientifically valid and practical and are increasingly used in Europe and worldwide (Coquette et al., 2003; do Nascimento Pedrosa et al., 2017; Poumay et al., 2004). Unfortunately, the elimination of all animal-originated (including human-originated) products from the cell culture media used to generate and validate RHE and full-thickness skin models used for the testing of cosmetic ingredients has not been reported yet in the peer-reviewed literature.

To create epidermal substitutes such as RHE, the availability of primary keratinocytes or immortalized keratinocytes capable of keratinization, in addition to appropriate basal media, their growth supplements and adequate porous membranes for tissue growth at the air-liquid interface are all critical (Rosdy & Clauss, 1990). The COVID-19 pandemic, unfortunately, provoked sporadic interruptions in the supply chain of essential products, including the HKGS required to support epidermal reconstruction when using EpiLife medium. Consequently, many laboratories around the world relying on HKGS were impeded to reconstruct the epidermis. Seeking an alternative to HKGS, we identified cdAOF supplements provided by AvantBio corporation (HKSdaFREE + HKGE and HFSdaFREE) as an interesting option for generating RHE models for evaluation and acceptance under OECD cosmetic testing guideline 431 (OECD, 2015). Additionally, cdAOF supplements not only address the expectations of cosmetic testing’s ethical and marketing advantages (e.g. being animal cruelty-free), but also the advantages of a safer and diminished transmissible disease risk for reconstructive and regenerative medicine (Baltazar et al., 2023).

In our current study, we compare the standard BPE-containing HKGS supplement previously described for RHE production (Dijkhoff et al., 2021; Hall et al., 2022; Lemper et al., 2014) to the cdAOF supplements HKSdaFREE, HKSdaFREE + HKGE and the fibroblast-specific HFSdaFREE (Cook P et al., 2012; Cook PW et al., 2012). Additionally, to obtain a broad, cell-specific performance comparison on all the tested supplements, RHEs were produced from primary HEKn, primary HEKa and immortalized N/TERT keratinocytes (Dickson et al., 2000).

## MATERIALS AND METHODS

### Origin of cells

HEKa were isolated from abdominoplasty (Clinique St Luc, Namur, Belgium), as described (Minner et al., 2010). Samples from patients were obtained after written informed consent and in agreement with the principles and guidelines of the Declaration of Helsinki. HEKn were purchased from ThermoFisher Scientific (Cat#C0015C, Massachusetts, USA). Immortalized N/TERT keratinocytes were obtained from the J. Rheinwald laboratory (Harvard Medical School, Boston, USA) (Dickson et al., 2000).

### Culture condition and skin reconstruction

HEKa, HEKn and N/TERT keratinocytes were routinely cultured in EpiLife medium (ThermoFisher Scientific Cat#MEPI500CA, Massachusetts, USA) supplemented with HKGS (ThermoFisher Scientific Cat#S0015, Massachusetts, USA), penicillin 50 U/ml and streptomycin 50 µg/ml (Millipore Sigma, Cat#P4333, Missouri, USA) until reaching 70-80% of confluency. Keratinocytes were harvested, then seeded at density of 330,000 cells/cm² onto a polycarbonate filter with a pore size of 0.4 µm, as previously described (Poumay et al., 2004). To reconstruct the epidermis, the EpiLife medium with four different supplement formulations was used as follows: a) HKGS; b) HKSdaFREE (HKS), specific for human keratinocytes and corneal epithelial cells (AvantBio Inc., Cat#AVB01HKS, Lynnwood WA, USA); c) HKSdaFREE + HKGE (HKS + HKGE) (AvantBio Inc., Cat#AVB01HKS, Lynnwood WA, USA) and d) HFSdaFREE (HFS), specific for human fibroblasts (AvantBio Inc., Cat#AVB02HFS, Lynnwood WA, USA). All media used for the culture of RHEs received an additional 1.44 mM CaCl₂ to reach 1.5 mM Ca²⁺ final concentration. After 24 h of incubation at 37°C in a humidified atmosphere containing 5% CO₂, seeded cultures were exposed to the air-liquid interface. Media were additionally supplemented with 50 µg/ml vitamin C (Sigma-Aldrich Cat#49752, Saint Louis, MO, USA) and 10 ng/ml keratinocyte growth factor (R&D Systems Cat#251KG, Minneapolis, MN, USA). RHEs were cultured for ten additional days with media being changed every 2-3 days.

### Hematoxylin & eosin staining (H&E)

After 11 days of reconstruction, histological preparation of RHE and H&E staining of tissue sections were performed as previously described (E. De Vuyst et al., 2013). Briefly, the RHEs were fixed in 4% acetic formalin, dehydrated, and embedded in paraffin. 6 µm thick tissue sections were cut and placed onto microscope slides. After deparaffinization and rehydration, sections of RHE were stained with H&E, dehydrated and mounted with DPX (VWR International, Leuven, Belgium).

### Immunohistological staining

For immunolabeling, RHE sections were deparaffinized, rehydrated and rinsed with tap water. To detect specific proteins, the following primary antibodies were used: CK14 (dilution 1:50; Santa Cruz Cat#sc58724, CA, USA), CK10 (dilution 1:100; Dako Cat#M7002, Glostrup, Denmark), involucrin (dilution 1:200; Millipore Sigma Cat#I9018, Missouri, USA), loricrin (dilution 1:100; Abcam Cat#ab24722, Cambridge, UK), filaggrin (dilution 1:75; ThermoFisher Scientific Cat#M13440, Waltham, MA, USA) and Ki67 (dilution 1:50; Diagomics Cat#M3034, Blagnac Cedex, France). For immunofluorescence (IF) the secondary antibody used was Alexa 488-conjugated anti-mouse IgG (dilution 1:200; Invitrogen Cat#A11001, Carlsbad, CA, USA) or Alexa 488-conjugated anti-rabbit IgG (dilution 1:200; Invitrogen Cat#A11008, Carlsbad, CA, USA). For immunohistochemistry (IHC), the anti-rabbit Vectastain ABC HRP kit (dilution 1/200; Vector Laboratories Cat#PK4001, CA, USA) was used.

### Transepithelial electrical resistance measurement (TEER)

On the 11^th^ day of RHE culture, TEER was measured using Millicell® ERS-2 device (Millipore Corporation, Burlington, Massachusetts, USA).

### RNA extraction and Reverse Transcription-quantitative Polymerase Chain Reaction (RT-qPCR)

RNA isolation and RT-qPCR were performed as previously described (Progneaux et al., 2023). Briefly, RNA from RHE was extracted using the “ReliaPrep™ miRNA Cell and Tissue Miniprep System” (Promega Cat#Z6111, Wisconsin, USA) and its concentration measured by NanoDrop 1,000 spectrophotometer (ThermoFisher Scientific, Massachusetts, USA). cDNA was then synthesized using “SuperScript III RNase H-Reverse Transcriptase Kit” (Thermo Fisher Scientific Cat#18080093, Massachusetts, USA). SYBR Green method (Eurogentec, Liège, Belgium) and specifically designed primers (Eurogentec, Liège, Belgium; Millipore Sigma) listed in Table 1 were used. Gene expression is normalized to the expression of ribosomal protein P0 (RPLP0) as reference (housekeeping) gene (Minner et al., 2010). Fluorescence of SYBRgreen was measured using LightCycler 96 (Roche Diagnosis, Vilvoorde, Belgium).

## RESULTS

### The chemically defined animal origin-free supplements allow reconstruction of a functional human epidermis that displays similar, but lower TEER values, with basal layer keratinocytes exhibiting a less cylindrical morphology

In vitro 3D skin models were reconstructed using HEKn, HEKa or N/TERT keratinocytes. Each type of RHE was cultured with HKGS, HKS, HKS + HKGE or HFS supplement. To evaluate the morphology of RHE, H&E staining was performed, revealing that every RHE corresponded to fully differentiated epidermis covered with a typical keratinized stratum corneum (Fig. 1A). However, basal keratinocytes in models reconstructed with HKS, HKS + HKGE or HFS exhibited less frequent cylindrical cell morphology when compared to RHE produced with HKGS. The less cylindrical basal epidermis was most evident in the RHEs cultured with the fibroblast specific HFS supplement. Nonetheless, the fibroblast-specific cdAOF supplement still produced an RHE.

**Fig. 1.**
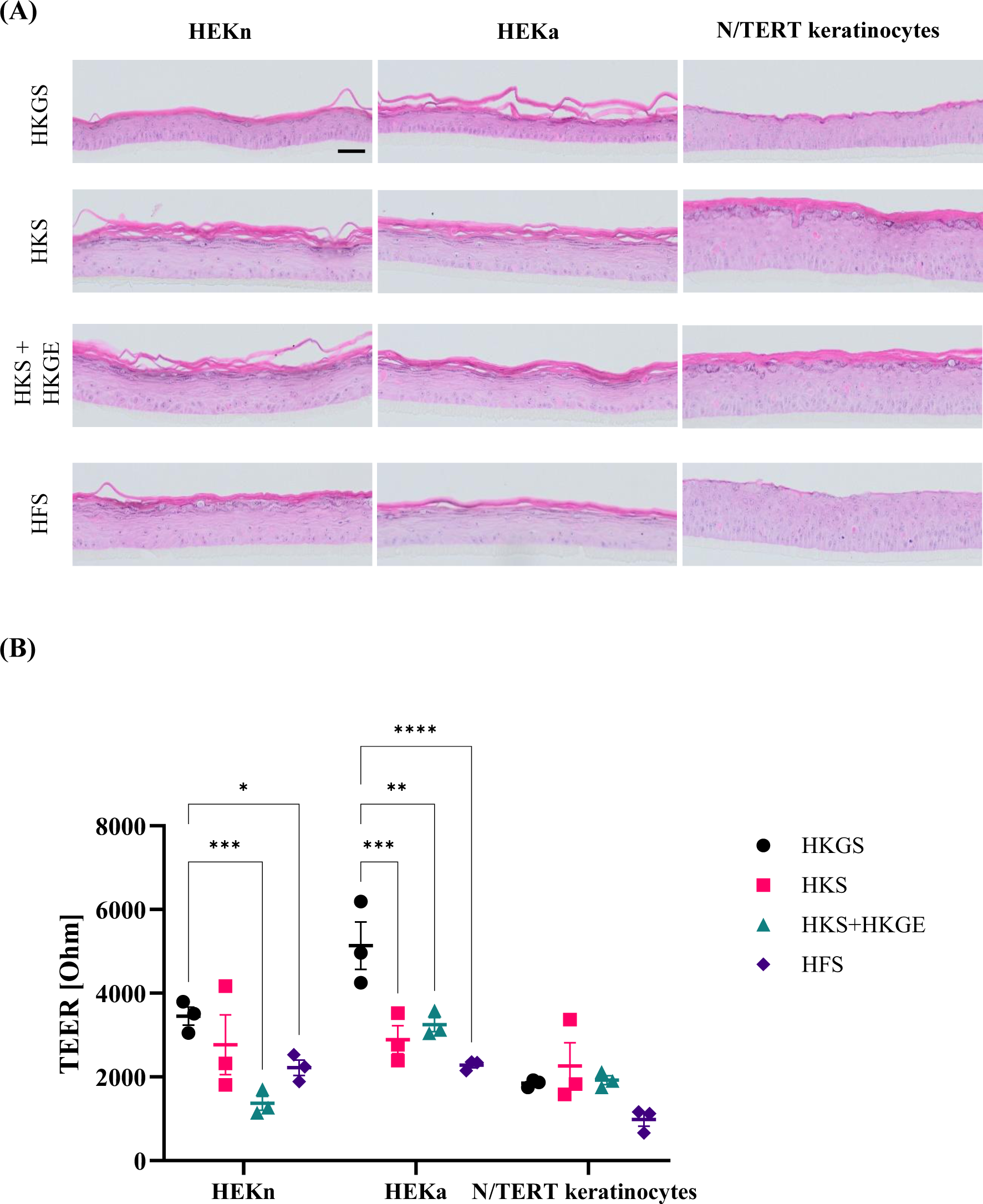
Microanatomy and barrier differences of the skin models reconstructed with HEKn, HEKa or N/TERT keratinocytes, as cultured with HKGS, HKS, HKS + HKGE or HFS supplement, respectively. **(A)** Histological sections of RHE stained with H&E. Scale bar: 50 µm; pictures are representatives of n = 3 experiments with a total of 3 skin models at each experimental condition. Magnification (x100) **(B)** Trans-epidermal electrical resistance (TEER) measurements performed on RHE (n = 3). Data represent mean ± SD. Two-way ANOVA was performed, followed by Dunett’s tests. *p<0.05, **p<0.01, ***p<0.001, ****p<.0001.

The barrier function of RHE was evaluated by TEER measurement (Fig. 1B). First, average TEER values of 1,500 to 3,000 Ohms were determined, respectively for HEKn- and HEKa-derived RHEs cultured in cdAOF HKS media, and revealed results close to those observed with RHE cultured in HKGS. Additionally, comparing HEKn and HEKa cultured in HKGS versus HKS, TEER values were higher in RHEs derived from HEKa cultured in HKGS, but there was no statistically significant difference in the TEER values from RHEs derived from HEKn. The barrier as measured by TEER appears to be generally higher when reconstructing epidermis from HEKa as compared to HEKn. Finally, we observed no significant difference in TEER values between any of the supplements on RHEs derived from immortalized N/TERT keratinocytes.

### Assessment of cellular proliferation via Ki67 expression in RHEs derived from HEKn, HEKa or N/TERT keratinocytes

The proliferative activity was evaluated using Ki67 marker-specific IHC staining of the RHEs (Fig. 2A). From the results, cell proliferation not only depends on the supplement used for epidermal reconstruction, but it also varies from one type of keratinocyte to another. Notably, keratinocytes obtained from a neonatal donor (HEKn) exhibited the highest frequency of Ki67 positive cells, along the basal layer, particularly when utilizing the HKGS supplement (Fig. 2B). Additionally, cdAOF supplements demonstrate an increased impact on the proliferation of N/TERT keratinocytes as compared to the animal-originated HKGS supplement.

**Fig. 2.**
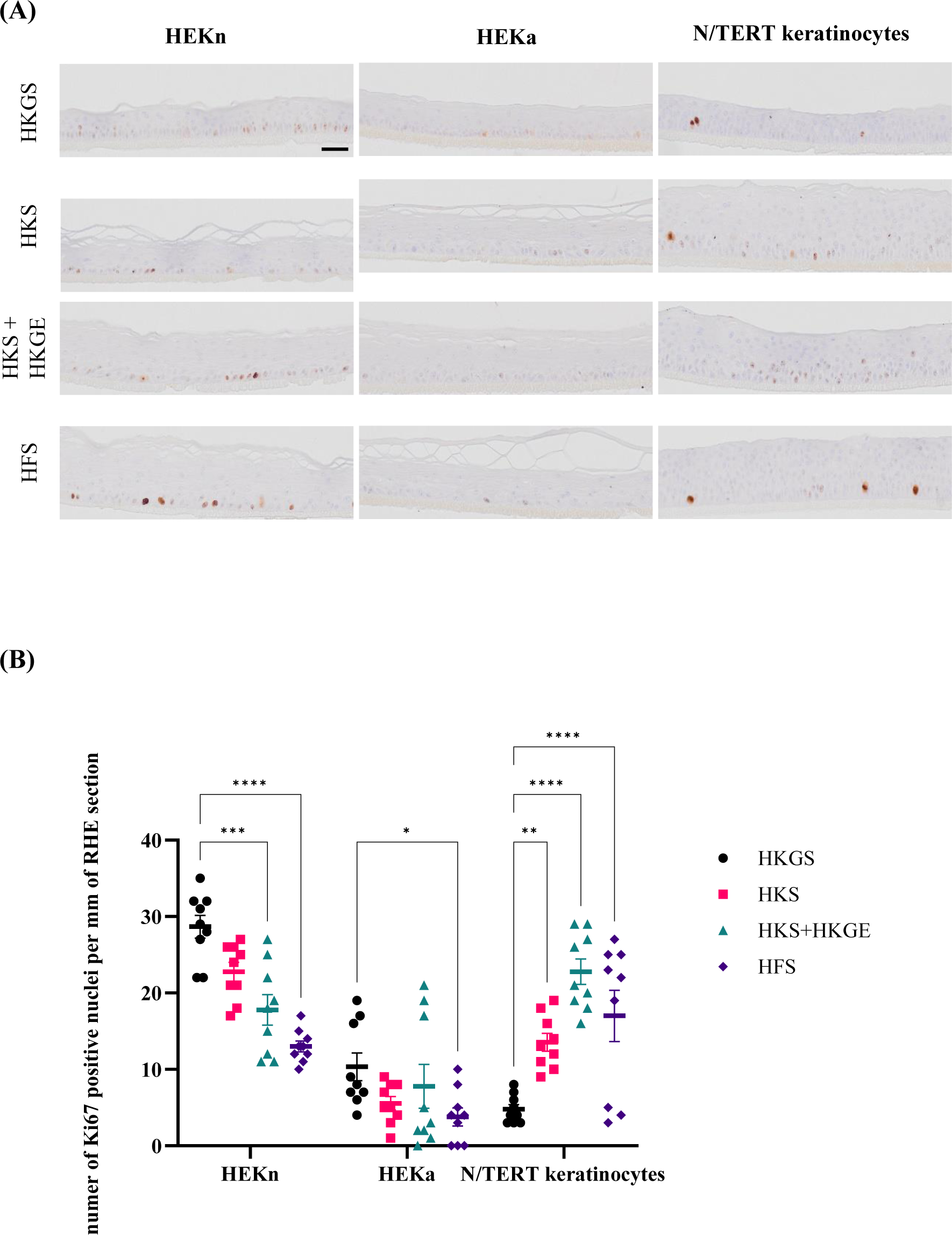
Proliferation of keratinocytes in RHE via Ki67 immunolocalization. **(A)** IHC staining for Ki67 revealed differences in proliferation marker intensity in skin models reconstructed with HEKn, HEKa or N/TERT keratinocytes cultured with HKGS, HKS, HKS + HKGE or HFS supplements (n = 3). Scale bar: 50 µm. Magnification (x100). **(B)** Quantification of proliferation marker Ki67 by double-blind test (n = 3). Data represent mean ± SD. Two-way ANOVA was performed, followed by Dunett’s tests. *p<0.05, **p<0.01, ***p<0.001, ****p<0.0001.

### Differentiation marker immunolocalization in RHEs derived from HEKn, HEKa or N/TERT keratinocytes

To assess differentiation-related endpoints after epidermal reconstruction, immunostaining of several differentiation markers was performed. Immunostaining indicated that RHE produced from HEKn more strongly expresses cytokeratin 14 (CK14), a marker of basal keratinocytes, when compared to HEKa or N/TERT keratinocytes (Fig. 3A). Interestingly, the expression of CK14 was not directly related to the type of cell culture supplement used for tissue reconstruction. Immunolabeling of involucrin (IVL) displayed elevated and premature basal expression when RHE were cultured with the cdAOF supplements HKS, HKS + HKGE or HFS, as compared to RHE cultured in HKGS (Fig. 3B). Immunofluorescent staining of CK10 (Fig. 3C), LOR (Fig. 3D), or FLG (Fig. 3E) appear similar in RHE produced with cdAOF culture supplements and in RHE cultured with animal originated products (HKGS) and normal human skin (not shown).

**Fig. 3.**
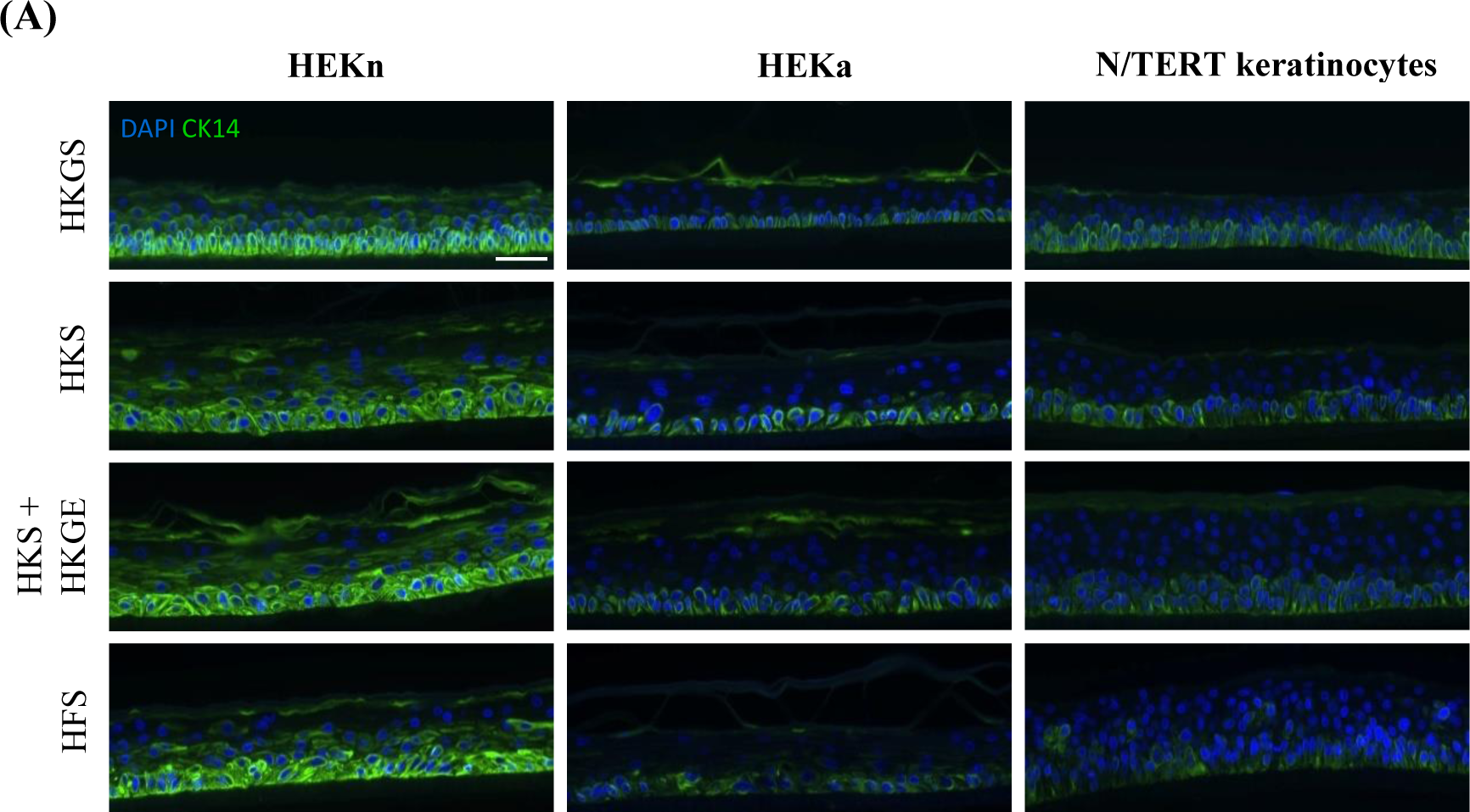
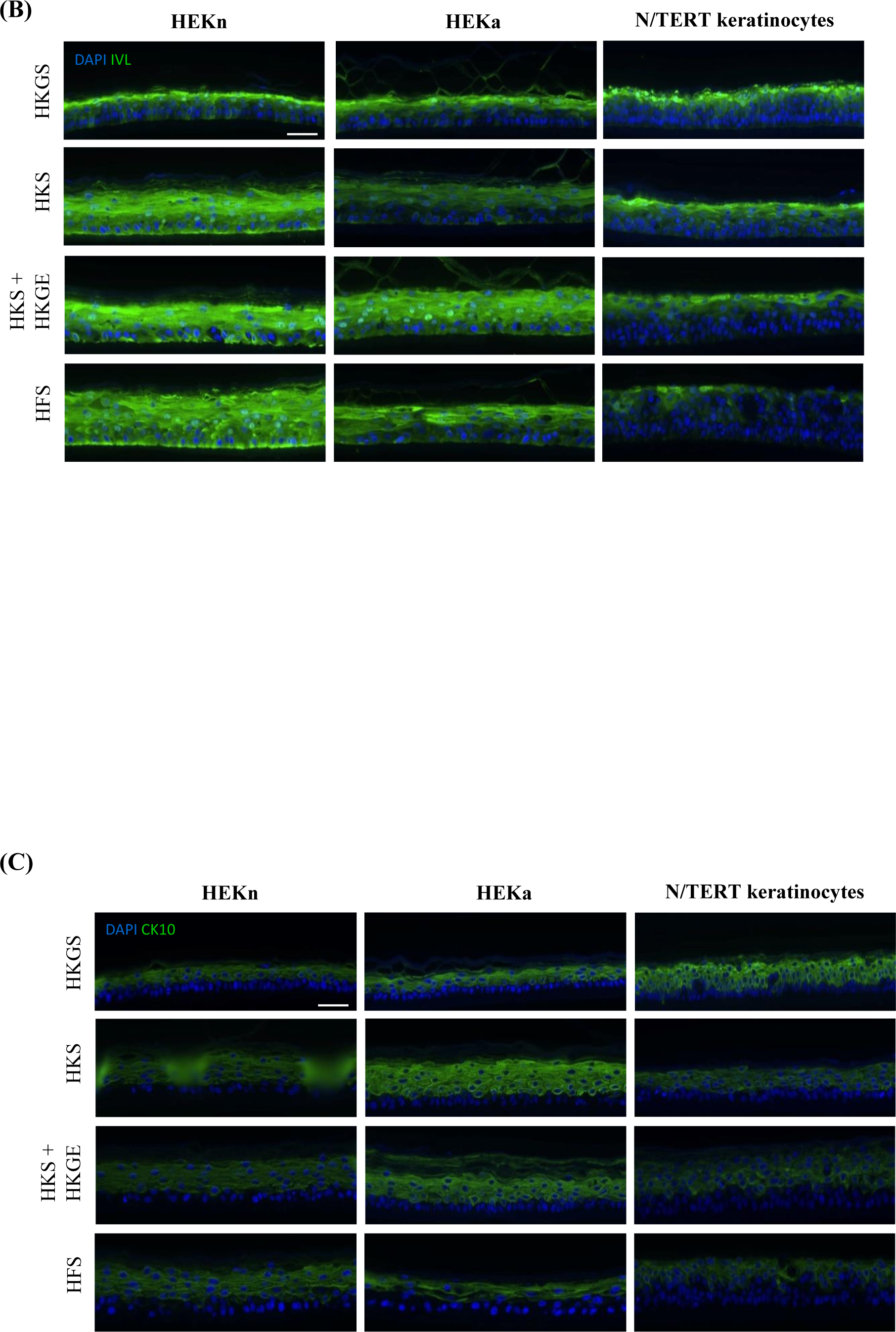
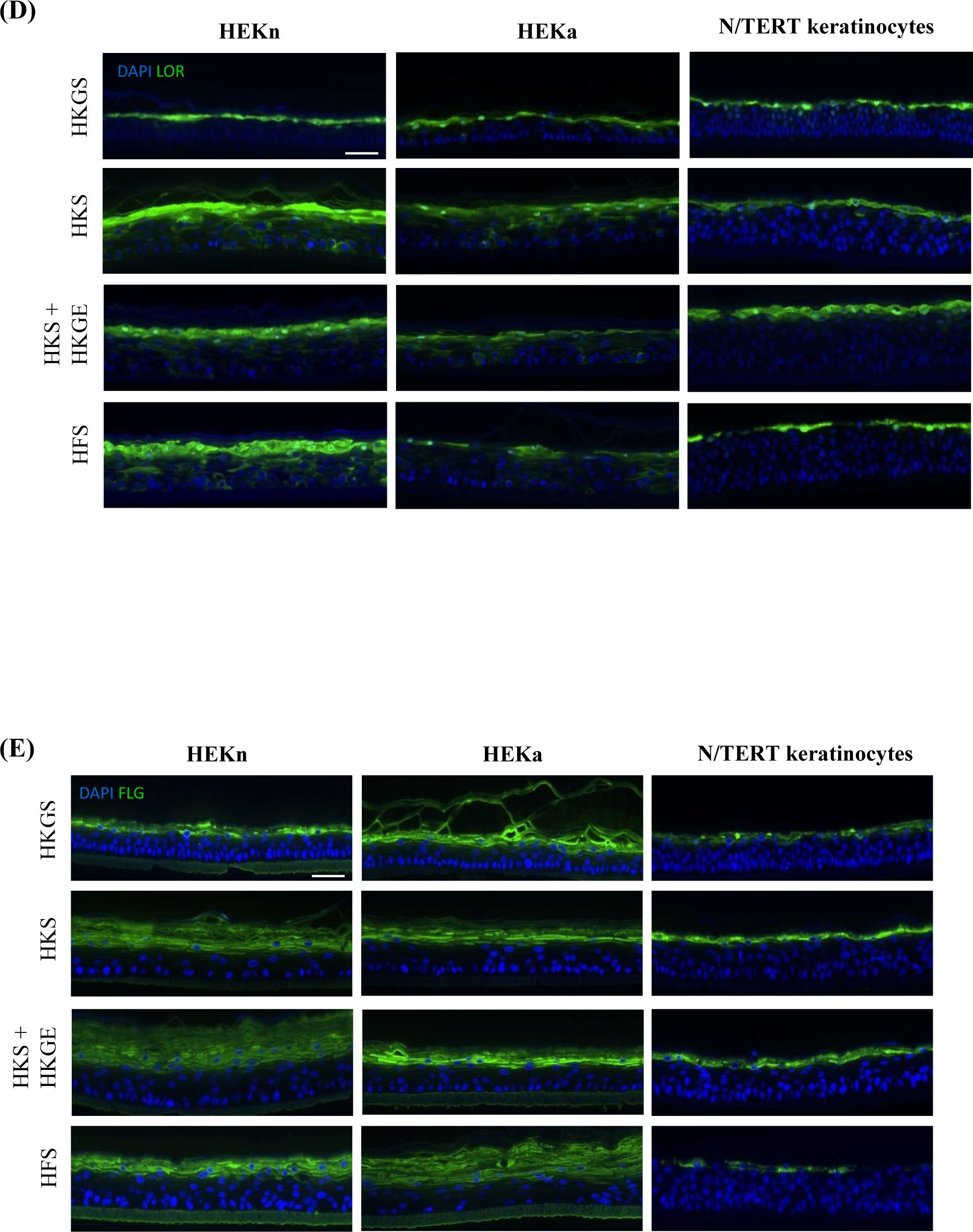
Immunostaining and localization of differentiation-related markers in RHE. **(A)** Cytokeratin 14 (CK14). **(B)** Involucrin (IVL). **(C)** Cytokeratin 10 (CK10) **(D)** Loricrin (LOR). **(E)** Filaggrin (FLG). Nuclei were stained with DAPI (blue). Scale bar: 50 µm. Pictures are representatives of 3 experiments with a total of 3 skin models for each experimental condition.

### Elevated levels of mRNA for interleukin 1-alpha (IL-1α) and other markers indicate an inflammatory-proliferative gene expression profile in RHEs cultured under cdAOF conditions

RHEs reconstructed from HEKn using the HKS supplements exhibited elevated levels of interleukin 1-alpha (IL-1α), amphiregulin (AREG), heparin-binding EGF-like growth factor (HB-EGF) and hyaluronan synthase 3 (HAS3) mRNA when compared to RHE cultured in HKGS. Similarly, RHE reconstructed from HEKn in the fibroblast-dedicated HFS supplement displayed elevated levels of IL-1α and AREG mRNA. The analysis of transcripts also demonstrated that HAS3, AREG and HB-EGF, three markers linked to inflammation, autocrine proliferation, inflammatory-proliferative skin pathologies such as psoriasis (Cook PW et al., 2004b) or exposure of keratinocytes to Th2 cytokines (É. De Vuyst et al., 2018) were significantly increased in RHE produced with HEKn cultured in HKS or HKS + HKGE supplements. In contrast, differentiation markers in RHE derived from HEKn cultured in HKS displayed significant changes in FLG and IVL mRNA levels. CK14, CK10, LOR and IL-8 gene expression appeared similar in RHE derived from HEKn cultured in the cdAOF supplements and in RHE derived from HEKn cultured in HKGS.

RHE from HEKa cultured in HKS + HKGE also demonstrated upregulated levels of IL-1α mRNA (Fig. 4B). Culturing RHE with the HKS + HKGE supplement associated with a significant increase in expression of CK10 and IVL transcripts (Fig. 4B). RHE derived from N/TERT keratinocytes culture in HKS or HKS + HKGE produce significantly elevated expression of IL-1α and AREG (Fig. 4C). Finally, our data reveal no significant changes in the expression of IL-8, under any condition.

**Fig. 4.**
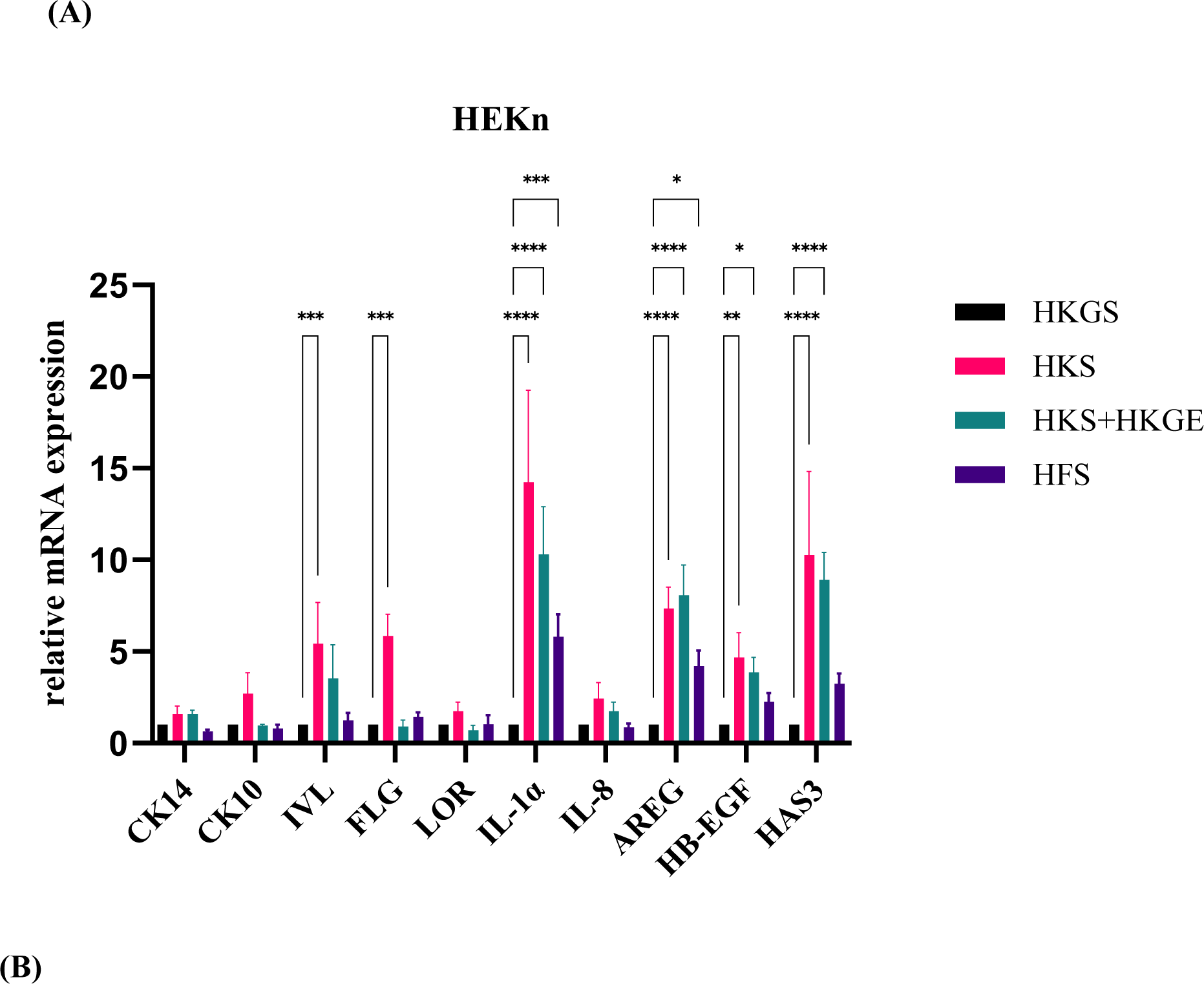
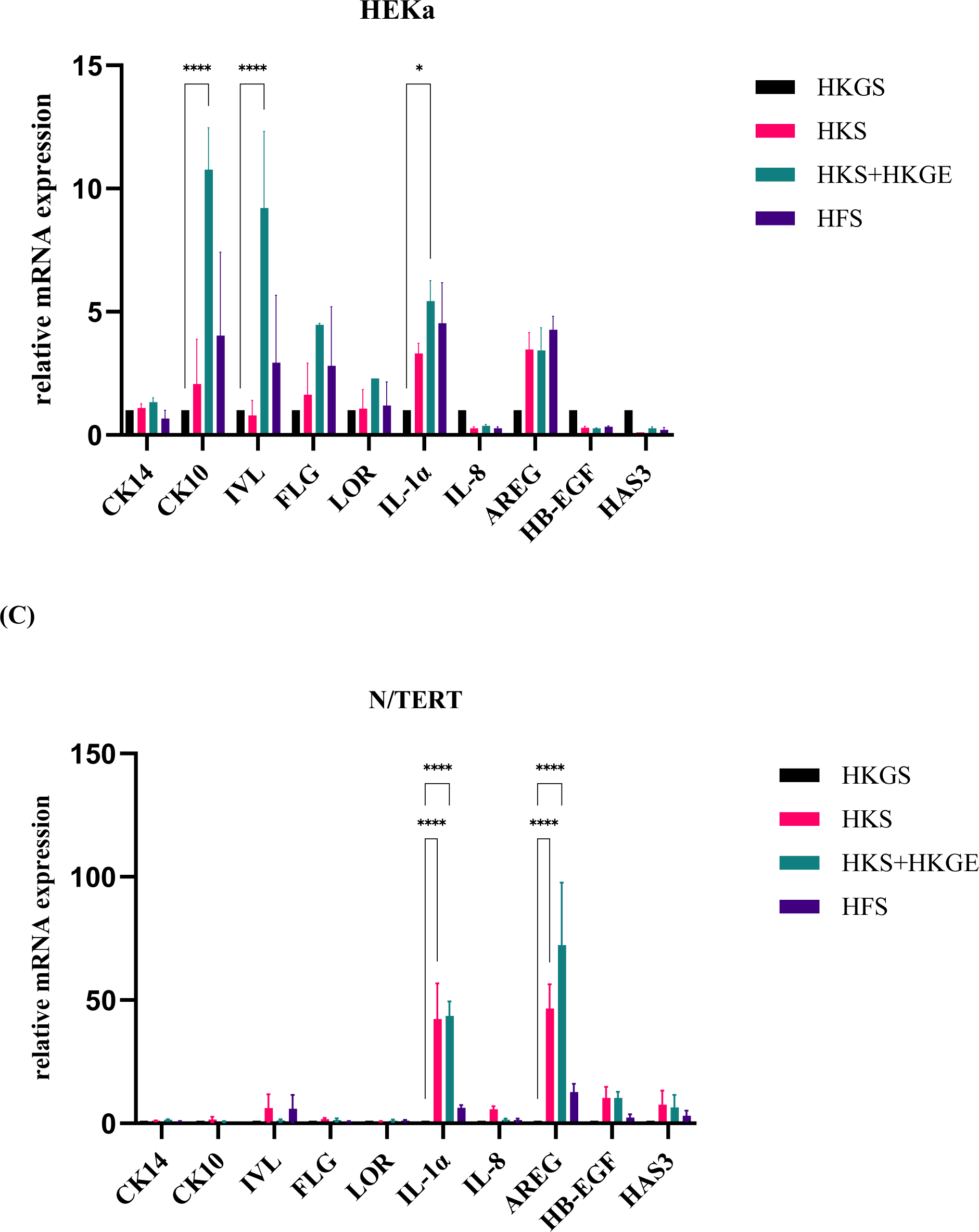
Transcriptional study of genes related to epidermal differentiation, inflammation and proliferation. mRNA expression was assessed by RT-qPCR. Gene expression in RHE constructed of **(A)** HEKn, **(B)** HEKa and **(C)** N/TERT keratinocytes. Data represent mean ± SEM (n = 3). Two-way ANOVA test was performed, followed by Dunnett’s tests. *p<0.05, **p<0.01, ***p<0.001, ****p<0.0001.

## DISCUSSION & CONCLUSIONS

Reducing the direct use of animals as well as the use of cell culture products derived from animals or humans is an inevitable progression for conducting research in accordance with the 3Rs principle (Díaz et al., 2021). Additionally, this progression reduces the risk of adverse inflammatory reactions or infectious events in reconstructive and regenerative cutaneous therapies. Over time, reconstructed skin models have demonstrated significant improvement with respect to accuracy and reliability. They are designed to mimic the structure and function of human skin, making them suitable substitutes for animal testing (Duval et al., 2003; Flaten et al., 2015; Kandárová et al., 2006). RHE are the simplest model dedicated to analyze some epidermal properties in vitro (Poumay & Faway, 2023). However, the production of RHE models generally relies on the use of HKGS or other animal-originated (including human-originated) components. This raises concerns regarding not only animal cruelty for those cell culture systems using reagents such as bovine-derived BPE and FBS, but also safety concerns when animal and human components exposed RHE tissue are used for reconstructive and regenerative medicine applications.

Additionally, considerable problems with the supply of the HKGS product during COVID-19 pandemic occurred, stagnating the research of many scientists in the RHE research field (Poumay & Faway, 2023). To test new alternatives and obtain a wider perspective regarding the use of cdAOF supplements for RHE reconstruction, we tested three different types of human keratinocytes comparing HKGS against the cdAOF supplements. We note that RHE derived from N/TERT keratinocytes, or similar immortalized keratinocyte cell lines, might reduce the cost of RHE production (Smits et al., 2017) and allow for easier genetic manipulation through techniques that include CRISPR-Cas DNA-editing (Evrard et al., 2021; Progneaux et al., 2023).

We demonstrate that all the cdAOF supplements produce functional RHE from either HEKn, HEKa or N/TERT keratinocytes. RHE produced with cdAOF supplements exhibit all the typical features of human skin morphology with respect to stratification of the basal, spinous, and granular layers, leading to a well-keratinized stratum corneum. Notably, the stratum corneum layer is thinner in RHEs produced from N/TERT keratinocytes but is nonetheless similar in thickness to what is observed in epidermis reconstructed with N/TERT keratinocytes in HKGS.

Nonetheless, there are some differences when comparing RHE produced using cdAOF supplements with RHE produced using HKGS. Basal keratinocytes of the human epidermis normally exhibit cylindrical shape (Yousef et al., 2023). However, in the current study, the animal-free cdAOF supplements produced RHEs displaying a less cylindrical basal layer, especially when HFS was used as an alternative to HKGS. On the other hand, HFS still promoted epidermal reconstruction, even though HFS was developed specifically for dermal fibroblast culture, rather than for epidermal keratinocyte culture. IVL is a late differentiation marker in human skin and normally localizes in the upper spinous and granular layers (Furue, 2020; Kanitakis et al., 1987). Using immunofluorescence to study involucrin localization in RHE, expression was found in the basal layer of RHE cultured using cdAOF supplements, much earlier than normally observed in human skin and other RHE models (Fig. 3B). Premature, enhanced expression of involucrin occurs in some skin disorders linked to imbalance in the keratinization process (Kanitakis et al., 1987). For instance, a similar premature distribution of involucrin was found in RHE derived from the keratinocytes of a patient suffering Darier disease (Lambert de Rouvroit et al., 2013). Conversely, immunoreactivity of both CK14, which is a marker of proliferative keratinocytes in the basal layer (Vaidya & Kanojia, 2007) and CK10, a marker of early differentiation in the suprabasal layers, appear correctly localized in RHE cultured with cdAOF supplements (Fig. 3). Similarly, other markers of late epidermal differentiation, such as LOR and FLG, are normally expressed in the granular layer of cdAOF RHEs, bringing us to conclude that all the cdAOF supplements conferred production of a functional RHE. This is also supported by the fact that HKS + HKGE was utilized as a novel cdAOF epidermal cell bioink to generate a xenofree, fully vascularized, full-thickness 3D-bioprinted skin equivalent successfully engrafted onto mice (Baltazar et al., 2023).

Our current study further revealed that RHE derived from N/TERT keratinocytes (Fig. 2B) exhibited increased proliferation rates when using cdAOF supplements as compared to the animal product-containing HKGS supplement. Primary keratinocytes in RHE derived from HEKn and HEKa show greater proliferation rates when cultured with HKGS containing animal-originated products (BPE), than with cdAOF supplements. Interestingly, our study also suggests that frequency of Ki67-positive cells is closely related to the type of keratinocytes used to produce the RHE. Still, TEER analysis revealed that barrier formation is more complete when HKGS is used to generate an RHE, as compared to the use of the cdAOF supplements. RT-qPCR results (Fig. 4) reveal that models reconstructed with cdAOF supplements exhibit signs of an inflammatory-proliferative response since increased levels of IL-1α, HA3, HB-EGF and AREG mRNA are present. Since the composition of AvantBio cdAOF supplements are proprietary, identification of specific reagents included in the HKS + HKGE formulation that are responsible for the inflammatory-proliferative gene expression profile as well as the premature expression of involucrin and somewhat lower TEER values are currently under investigation.

In summary, RHE produced with cdAOF supplements such as HKSdaFREE or HKSdaFREE + HKGE, accurately mimic almost all the structural and functional properties (e.g. differential gene expression, stratification, and similar TEER values) of the human epidermis, despite premature and elevated involucrin expression es well as fewer columnar basal keratinocytes. The use of cdAOF cell culture supplements for epidermal reconstruction marks significant progress in the field of animal cruelty-free cosmetics, cosmeceuticals and aesthetic medicine, and safer regenerative medicine as well. This highlights the need for more advanced animal origin-free biomedical research and development. Moreover, the availability of cdAOF cell culture systems provides alternatives to the supply chain interruption of conventional animal product-containing cell culture supplements. Finally, the cdAOF approach not only aligns with ethical considerations to reduce the general use of animal-originated products, but also offers other compelling advantages, such as reducing the risk of transmission of both human-originated and zoonotic diseases during regenerative medical procedures.

## SUPPLEMENTARY TABLES

**Tab. 1.**
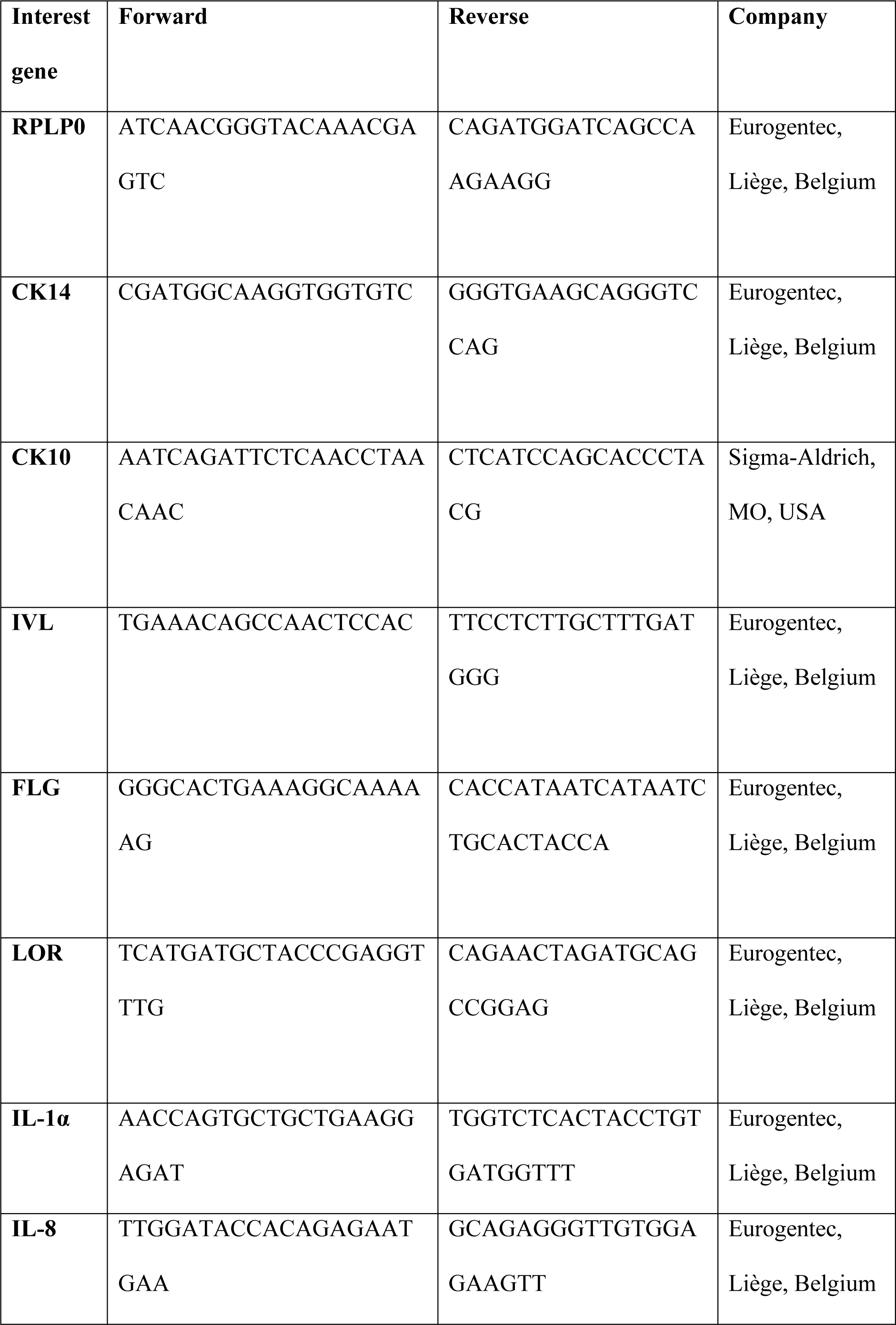
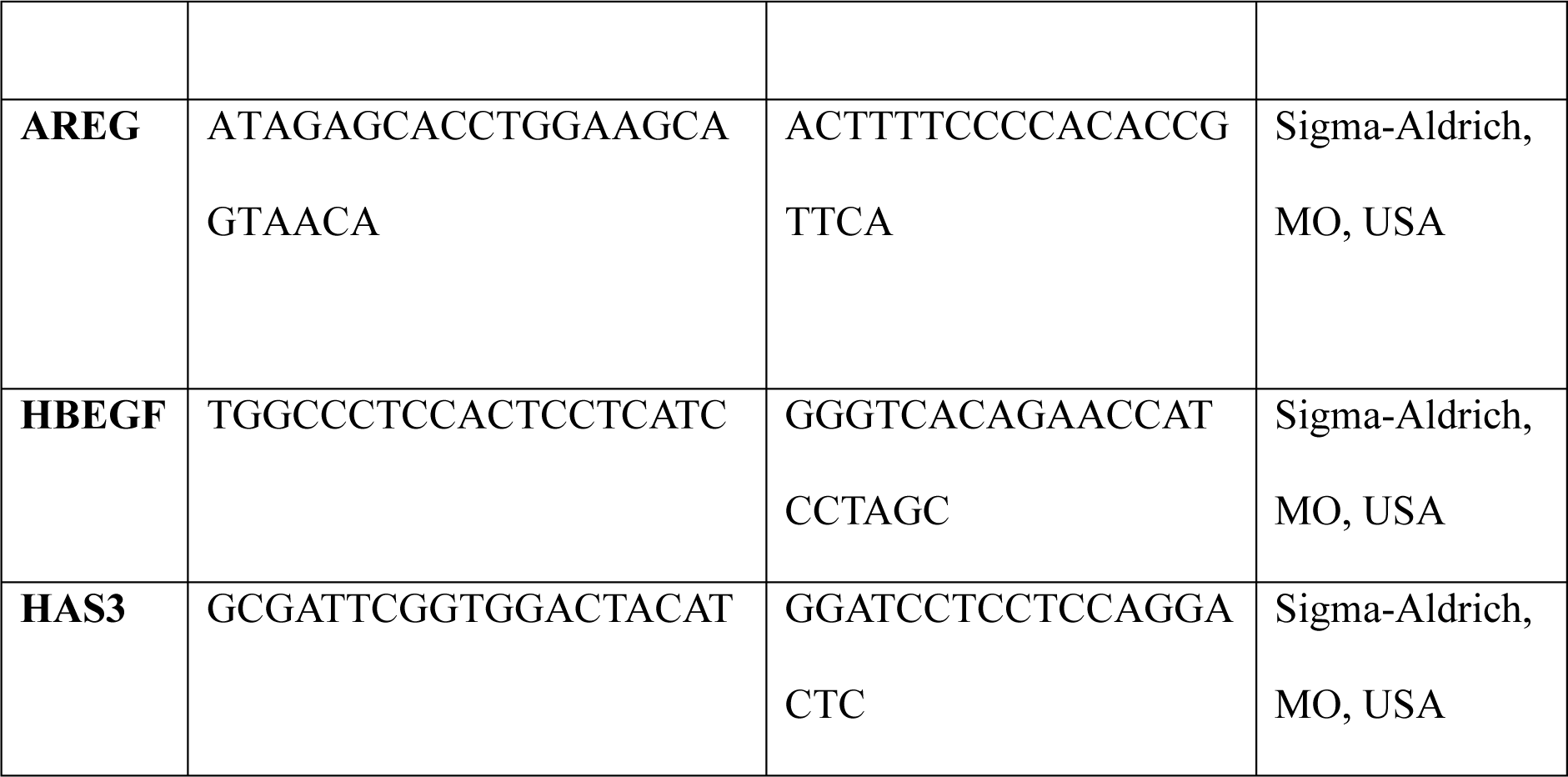
Sequence of primers used for RT-qPCR.

